# A Single-shot ChAd3 Vaccine Provides Protection from Intramuscular and Aerosol Sudan Virus Exposure

**DOI:** 10.1101/2024.02.07.579118

**Authors:** Anna N. Honko, Ruth Hunegnaw, Juan I. Moliva, Aurélie Ploquin, Caitlyn N M Dulan, Tamar Murray, Derick Carr, Kathryn E. Foulds, Joan B. Geisbert, Thomas W. Geisbert, Joshua C. Johnson, Suzanne E. Wollen-Roberts, John C. Trefry, Daphne A. Stanley, Nancy J. Sullivan

## Abstract

Infection with Sudan virus (SUDV) is characterized by an aggressive disease course with case fatality rates between 40-100% and no approved vaccines or therapeutics. SUDV causes sporadic outbreaks in sub-Saharan Africa, including a recent outbreak in Uganda which has resulted in over 100 confirmed cases in one month. Prior vaccine and therapeutic efforts have historically prioritized Ebola virus (EBOV), leading to a significant gap in available treatments. Two vaccines, Erbevo^®^ and Zabdeno^®^/Mvabea^®^, are licensed for use against EBOV but are ineffective against SUDV. Recombinant adenovirus vector vaccines have been shown to be safe and effective against filoviruses, but efficacy depends on having low seroprevalence to the vector in the target human population. For this reason, and because of an excellent safety and immunogenicity profile, ChAd3 was selected as a superior vaccine vector. Here, a ChAd3 vaccine expressing the SUDV glycoprotein (GP) was evaluated for immunogenicity and efficacy in nonhuman primates. We demonstrate that a single dose of ChAd3-SUDV confers acute and durable protection against lethal SUDV challenge with a strong correlation between the SUDV GP-specific antibody titers and survival outcome. Additionally, we show that a bivalent ChAd3 vaccine encoding the GP from both EBOV and SUDV protects against both parenteral and aerosol lethal SUDV challenge. Our data indicate that the ChAd3-SUDV vaccine is a suitable candidate for a prophylactic vaccination strategy in regions at high risk of filovirus outbreaks.

**One Sentence Summary:** A single-dose of ChAd3 vaccine protected macaques from lethal challenge with Sudan virus (SUDV) by parenteral and aerosol routes of exposure.

## INTRODUCTION

Sudan virus (SUDV) is a filovirus that causes Ebola virus disease (EVD) in humans with case fatality rates between 40-100% (*1*). The first documented outbreak of SUDV occurred in 1976 in present-day South Sudan(*1*). Since then, SUDV has continued to cause sporadic outbreaks in sub-Saharan Africa with the latest occurring in 2022 in Uganda(*2*). The sporadic re-occurrence of EVD outbreaks suggests that the likely cause is zoonotic in nature, although EVD outbreaks have also been attributed to person-to-person transmission from EVD survivors (*3, 4*). Even though two vaccines (Erbevo^®^ and Zabdeno^®^/Mvabea^®^) have been licensed for use against the related Ebola virus (EBOV), these countermeasures are infective against SUDV (*4, 5*). Thus, effective vaccines against SUDV are urgently needed.

Recombinant adenovirus vectors (rAd) have demonstrated effectiveness against filoviruses as a component of vaccine regimens, as shown for EBOV, which showed uniform (100% survival) protection in nonhuman primates (NHP) (*6–9*). A replication-defective chimpanzee adenovirus (ChAd) was chosen as an alternative viral vector because of its superior safety profile and low seroprevalence in humans, overcoming constraints of preexisting immunity to human adenovirus vectors (*10–14*). ChAd vectors have a long clinical history, having been used safely in more than 5,000 people throughout more than 13 Phase 1, Phase 2, and Phase 3 human clinical trials, including in more than 600 pediatric patients. These trials have established that ChAd vectors are safe and produced antibody titers that are equivalent to those correlated with NHP protection (*12, 15–25*). Furthermore, filovirus specific ChAd3 vaccines have shown efficacy as a vaccine for EBOV and Marburg virus (MARV) (*10, 26*), with rapid and durable protection.

Here, we demonstrate in a NHP animal model that a single dose of ChAd3 encoding the SUDV glycoprotein (GP) protects from lethal SUDV variant Boneface (SUDV/Bon) challenge at a 5 week interval. We also demonstrate that a single dose of monovalent ChAd3-SUDV provides full protection against SUDV/Bon up to six months and partial protection up to one-year post-immunization. We report a strong correlation between the level of GP-specific binding antibodies and survival outcome in NHP, which we identified as an immune correlate of protection. The antibody response against SUDV GP persisted for at least one year after immunization. Additionally, we show that a bivalent ChAd3 vaccine encoding the GPs from both EBOV and SUDV protected NHPs from both parenteral and aerosol lethal SUDV challenge. These findings, along with the safety and immunogenicity shown in humans, indicate the potential for ChAd3-vectored single-shot filovirus vaccines for rapid use in ring vaccination procedure to prevent EVD (*4, 5*). The safety, immunogenicity, and durability of ChAd3-SUDV make it an ideal candidate for future preparedness against SUDV outbreaks.

## RESULTS

### A single-shot ChAd3-SUDV vaccine uniformly protects against lethal SUDV challenge

ChAd3 vaccines confer uniform protection against lethal EBOV and MARV challenge five weeks after immunization (*10, 26*). Here, we evaluated efficacy of a replication-defective ChAd3-vectored vaccine for protection against SUDV in NHP at a dose found to be protective against EBOV and MARV (*10, 26*). Cynomolgus macaques were immunized intramuscularly (i.m.) with a single inoculation of 1×10^10^ PU of ChAd3 encoding SUDV GP (ChAd3-SUDV) and challenged with a uniformly lethal dose (1,000 PFU, i.m.) of SUDV/Bon 5 weeks later. The unvaccinated macaque succumbed to infection at day 12 post-challenge and exhibited a plasma viral load exceeding 1×10^7^ plaque forming units per milliliter (PFU/mL) (Fig. 1A and B). All vaccinated macaques survived the infection without detectable viremia, and no fever, weight loss or other clinical manifestations characteristic of EVD were observed (not shown). Together, these data demonstrate that a single-shot of ChAd3-SUDV vaccine at a dose of 1×10^10^ PU generates acute protective immunity against the lethal effects of SUDV infection.

**Figure 1.**
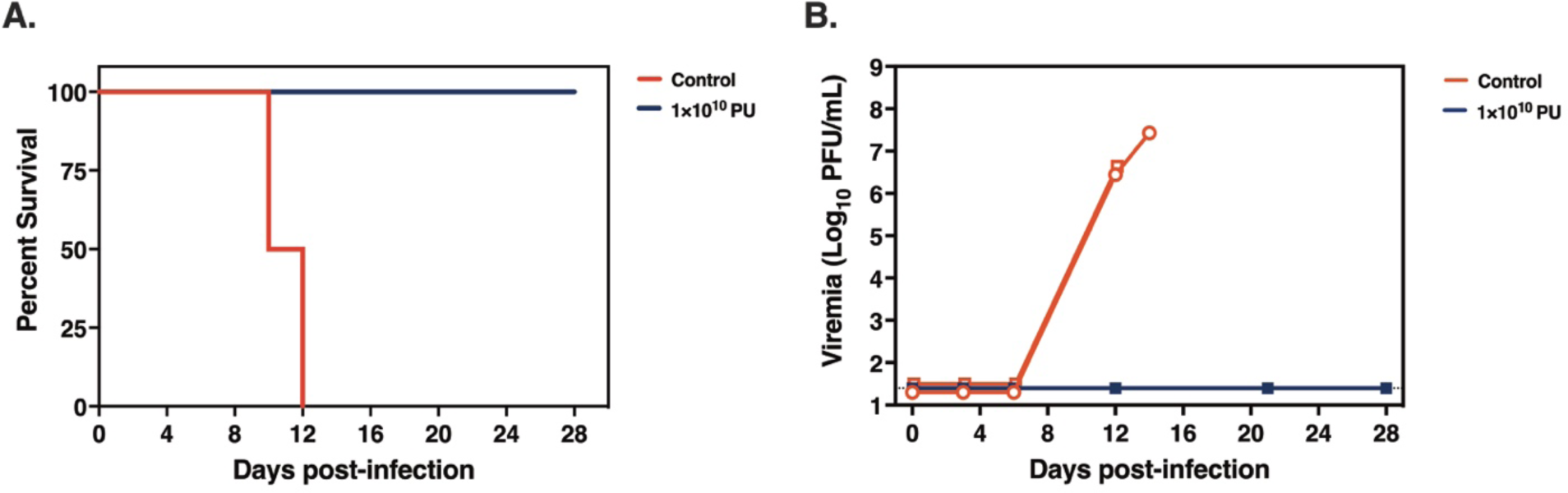
Single-shot ChAd3-SUDV vaccine uniformly protects against lethal SUDV challenge. Cynomolgus macaques were vaccinated with ChAd3-SUDV at a dose of 1×10^10^ PU via the i.m. route (*n*=5 total) or unvaccinated (*n*=2). Five weeks post-vaccination, NHP were challenged with a target dose of 1,000 PFU SUDV/Bon and followed for 28 days. (A) NHP survival following challenge (B) Plasma viremia by plaque assay in plaque-forming units per milliliter (PFU/mL). Open symbols indicate non-surviving animals.

### SUDV GP IgG titers predict protection

We next performed dose-ranging studies to define the minimal protective vaccine dose and to identify a candidate immune correlate of protection under conditions of infection breakthrough. Cohorts of cynomolgus macaques were vaccinated with ChAd3-SUDV at doses ranging between 1×10^7^ to 1×10^10^ PU. SUDV GP-specific antibody titers were measured four weeks later (Fig. 2A). Anti-SUDV GP titers and percent protection data from vaccine dose groups described in Figure 1: 1×10^10^ (*n =* 4) (Fig. 1A and B), were combined with data from the dose-ranging study for composite data representation across all doses in Figure 2 (Fig. 2). Both unvaccinated macaques succumbed after a lethal SUDV/Bon challenge. Macaques that received the 1×10^10^ PU vaccine dose uniformly survived without detectable viremia (Fig. 1A and B, Fig 2A and B), while mortality with viremia was observed in macaques that received doses below 1×10^10^ (Fig. 2B and C).

**Figure 2.**
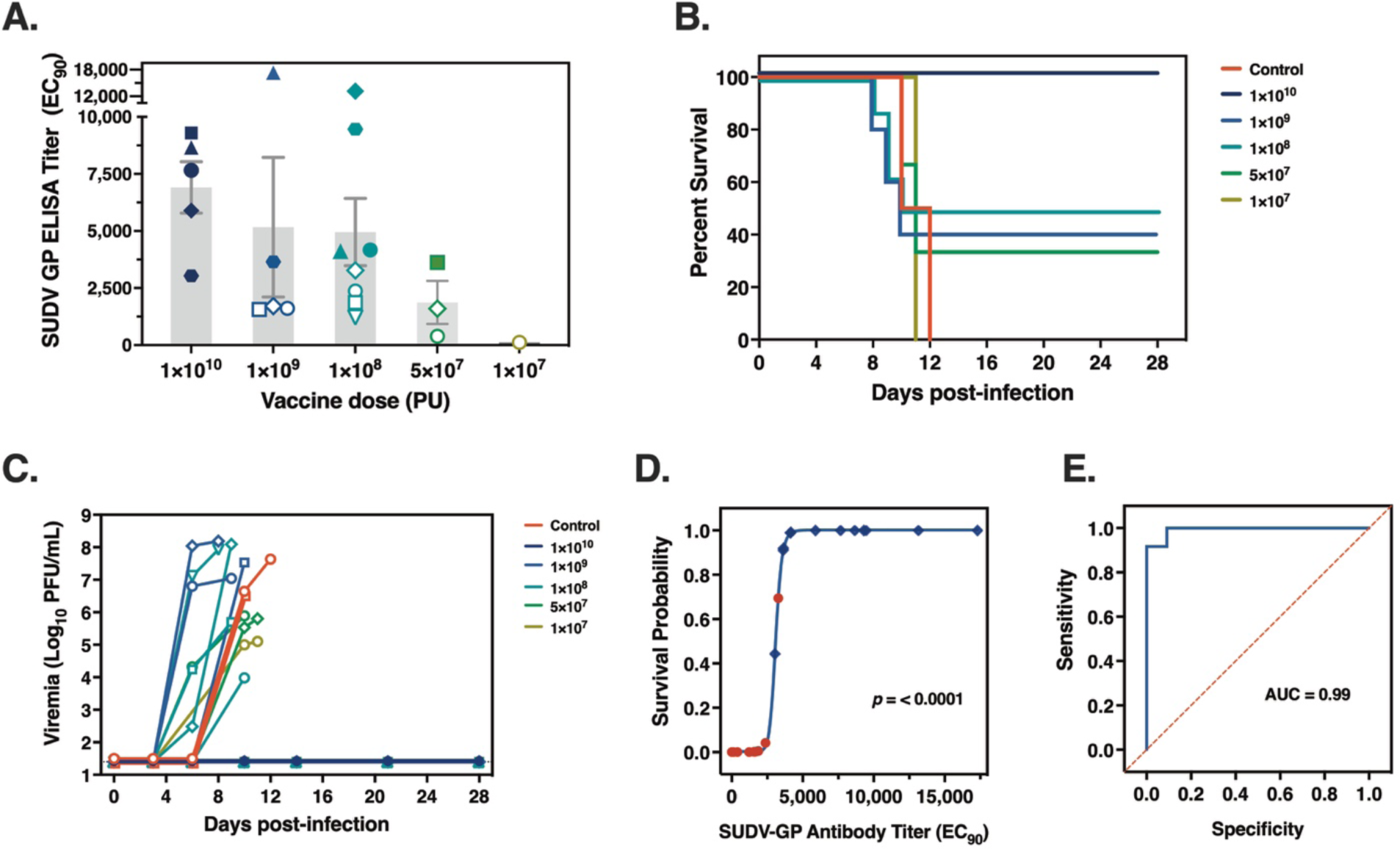
Anti-SUDV GP antibodies correlate with ChAd3-SUDV vaccine protection against lethal SUDV challenge. Groups of 5 cynomolgus macaques were vaccinated with ChAd3-SUDV at 1×10^10^, 1×10^9^, 1×10^8^, 5×10^7^ or 1×10^7^ PU via the i.m. route and challenged with 1,000 PFU SUDV/Bon five weeks later. (A) Plasma SUDV GP-specific ELISA titers at 4 weeks post-vaccination (EC_90_). Open/white symbols indicate non-surviving animals. (B) Percent survival following challenge. (C) Plasma viremia by plaque assay. (D) Univariate logistic regression of anti-SUDV GP EC_90_ titers and survival after lethal SUDV challenge. Regression was performed on ELISA IgG titers generated by ChAd3-SUDV vaccination of cynomolgus macaques (*n* = 22). Y-axis illustrates survival probabilities predicted by IgG titers on a continuous scale. Blue diamonds, survivors; red circles, non-survivors. Dotted lines: antibody titer associated with a survival probability of 85%. The p-value = two-sided Likelihood ratio test (chi-square) of the slope. (E) Receiver-operating characteristic (ROC) curve of data presented in panel D; AUC, area under the curve.

The average anti-GP EC_90_ titer generated in animals vaccinated at the uniformly protective vaccine dose of 1×10^10^ was 6909. Reduction of the vaccine dose to 1×10^9^ PU resulted in an average anti-GP EC_90_ titer of 5167, and infection breakthrough in 60% (3 out of 5) of the macaques. The 3 macaques that succumbed had the lowest titers of the group, averaging at an EC_90_ of 1628 compared to an average EC_90_ titer of 10475 for the two macaques that survived (Fig. 2A and B). A further reduction in dose to 1×10^8^ showed mean EC_90_ titers of 4954 and breakthrough in 50% (4 out of 8) of the macaques, which was comparable to the titers and breakthrough observed in response to the 1×10^9^ PU dose (EC_90_ 5167; 60%, respectively). The lowest vaccine doses of 5×10^7^ PU and 1×10^7^ resulted in a mean EC_90_ titer of 1432, significantly lower than titers obtained at the highest dose of 1×10^10^ PU (6909, *p* = 0.007). This reduction in dose was associated with a further increase in breakthrough to 75%, i.e. with 3 out of 4 macaques succumbing (Fig. 2A and B). Overall, 1×10^10^ PU ChAd3-SUDV was the minimum uniformly protective dose in these studies and significant reduction in antibody titers was associated with increased breakthrough.

Across individual animals in these studies, uniform protection (100%) was observed above a SUDV-GP ELISA EC_90_ titer of 3621. Mortality was 100% at a titer of 2371 and below. Between 3621 and 2371 one animal survived and one animal succumbed, indicating that the threshold for protection lies within this range. In summary, these data indicated that higher SUDV GP-specific antibody titers associated with protection.

Recently, we have reported that antibodies are a significant immune correlate of protection for ChAd3-MARV vaccine (*26*) which suggested that antibodies could be an immune correlate of protection for the ChAd-SUDV vaccine. Therefore, we tested the strength of the association between SUDV GP-specific titers and survival using univariate regression analysis with IgG titer as a continuous variable, and calculated survival probabilities over the range of antibody titers observed. A significant association was found between anti-SUDV GP titers and survival (*p* < 0.0001) indicating that quantity of anti-SUDV GP positively correlate with survival (Fig. 2D). Receiver-operating characteristic (ROC) analysis was performed to measure the sensitivity and specificity of anti-SUDV GP titers in predicting survival. An area under the curve (AUC) value representing the logistic model sensitivity and specificity (where the maximum is 1.0) of 0.99 was obtained, indicating that SUDV GP-specific antibody titers at 4 weeks post-vaccination provide robust discrimination between survivors and non-survivors. Therefore, the titer at 4 weeks post-vaccination is a sensitive predictor of acute protection induced by the ChAd3-SUDV vaccine (Fig. 2E).

### A single-shot ChAd3-SUDV vaccine provides durable protection from lethal SUDV challenge

In outbreak settings, a vaccine needs to provide rapid protection from the virus following immunization for effective control and utility in ring-vaccination protocols. However, an ideal vaccine will also provide extended immunity for protection throughout the duration of an outbreak response or for persons likely to come into contact with the virus in endemic regions. Having demonstrated the protective efficacy of ChAd3-SUDV vaccination at an acute timepoint of 5 weeks post-immunization, we next sought to evaluate the durable protection of the single-shot ChAd3-SUDV vaccine. Groups of macaques were immunized with 1×10^11^ PU of ChAd3-SUDV and their antibody titers were measured at weekly intervals prior to challenge at either 6 or 12 months post-vaccination. Complete protection from lethal challenge with SUDV/Bon was observed at 6 months, wherein vaccinated NHP exhibited no viremia throughout the study period (Fig 3B). Protection with ChAd3-SUDV extended to 12 months, where 3 out of 4 macaques survived. A single vaccinated animal succumbed at the 12 month timepoint. This animal had a pre-challenge EC_90_ titer of 941 at 48 weeks post-vaccination, where predicted survival was <10% per the logistic regression (Fig 3E). Transient viremia was detected in two of the survivors challenged at the 12 month timepoint, but in both cases resolved by day 10 post-challenge. Viremia did not appear to correlate with the pre-challenge SUDV-GP antibody titer, as these were the subjects with the lowest and the highest ELISA EC_90_ values (Fig 3D and 3F). Together, these findings demonstrate that a single-shot of 1×10^11^ PU ChAd3-SUDV provides protection against lethal infection for up to 1 year post-vaccination.

**Figure 3.**
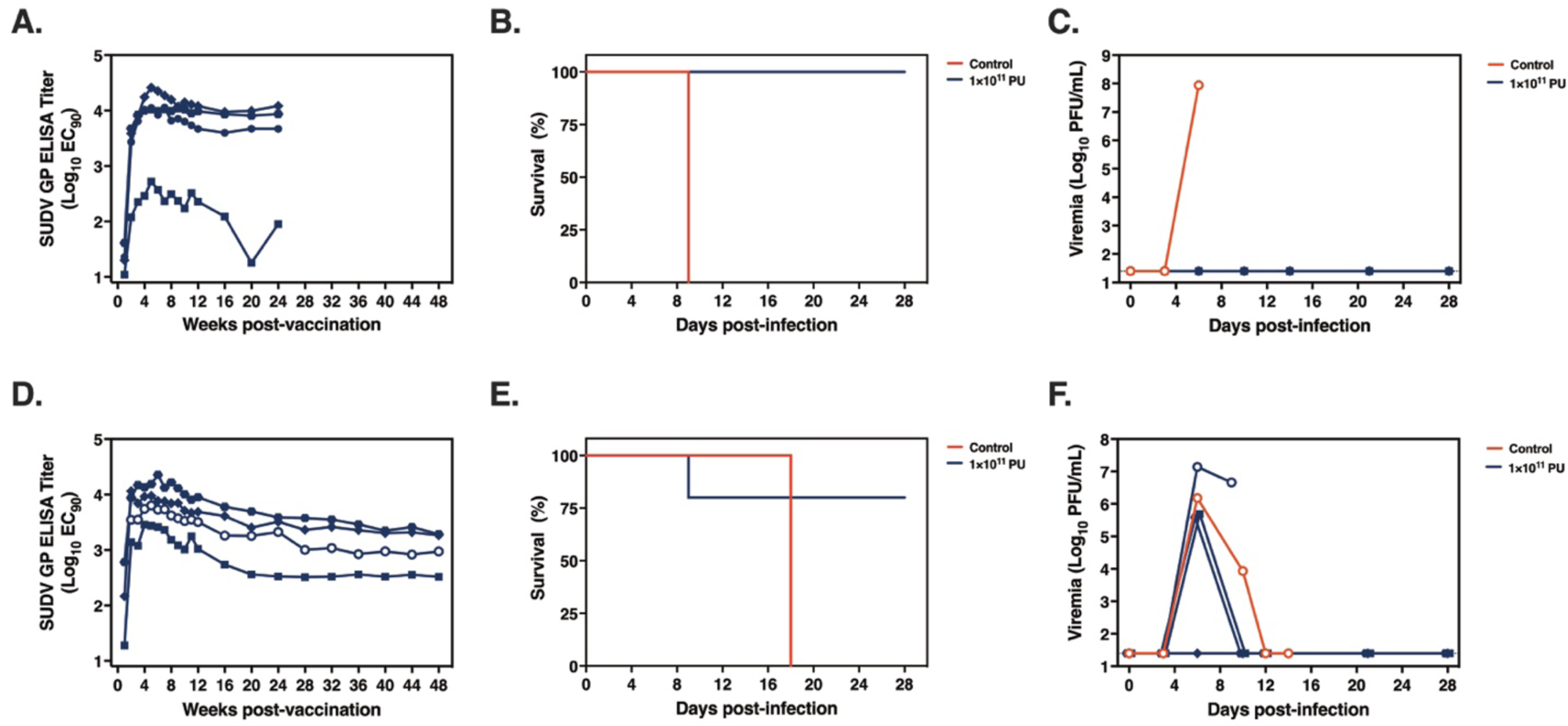
ChAd3-SUDV vaccine protects against SUDV challenge at durable timepoints. In two iterations, cynomolgus macaques (*n*=4) were vaccinated with 1×10^11^ PU ChAd3-SUDV via the i.m. route or unvaccinated (*n*=1). Plasma was collected at weekly intervals prior to i.m. challenge with 1,000 PFU SUDV/Bon at either 6-months (A-C) or 12 months (D-F). (A and B) Plasma SUDV GP-specific ELISA titers following immunization (EC_90_). Open symbols indicate non-surviving animals. (B and E) Survival following lethal SUDV challenge. (C and F) Circulating viremia as determined by plaque assay.

### A single-shot bivalent ChAd3-SUDV/ChAd3-EBOV vaccine uniformly protects against lethal SUDV challenge

SUDV and EBOV cause the most morbidity and mortality of the filoviruses and together have accounted for 38 of the 42 recorded outbreaks (*1*). In the context of a viral outbreak or epidemic, having countermeasures available that are effective across viral species would be an advantage. Towards this end, NHP vaccinated with a bivalent vaccine composed of equal ratios of ChAd3-SUDV and ChAd3 expressing the EBOV GP (ChAd3-EBOV) were evaluated in the lethal SUDV and EBOV challenge models. This bivalent vaccine has proven safe in humans (*11*), and the individual components of the bivalent vaccine proven effective in lethal NHP models (*10*). By evaluating the combined bivalent vaccine in lethal macaque models, we investigated whether the bivalent vaccine would remain as effective for protection as the individual component vaccines.

Groups of macaques were immunized with a single dose of equal ratios (*e.g.*, 1×10^10^ + 1×10^10^ PU) of bivalent ChAd3 (ChAd3-SUDV and ChAd3-EBOV) and challenged 5 weeks later with either SUDV/Bon (Fig 4) or EBOV variant Kikwit (EBOV/Kik) (Fig S1). The bivalent vaccine uniformly protected NHP from lethal EBOV exposure at a dose of 2×10^10^ PU. All of the SUDV challenged subjects immunized at 2×10^10^ PU survived, and only a single fatality was observed in the 2×10^9^ PU cohort. No detectable viremia was observed in the survivors. The SUDV-specific antibody titers at 4 weeks post-immunization were similar to those from equivalent monovalent-vaccinated animals (Fig. 2A). Additionally, we tested the bivalent vaccine in the SUDV/Gulu challenge model (Fig S1) where a dose of 2×10^11^ PU was uniformly protective with no viremia detected in any treated subject. This bivalent vaccine results in expanded breath of the vaccine beyond an individual viral species, and importantly, the addition of the second viral antigen did not reduce protective efficacy against SUDV.

**Figure 4.**
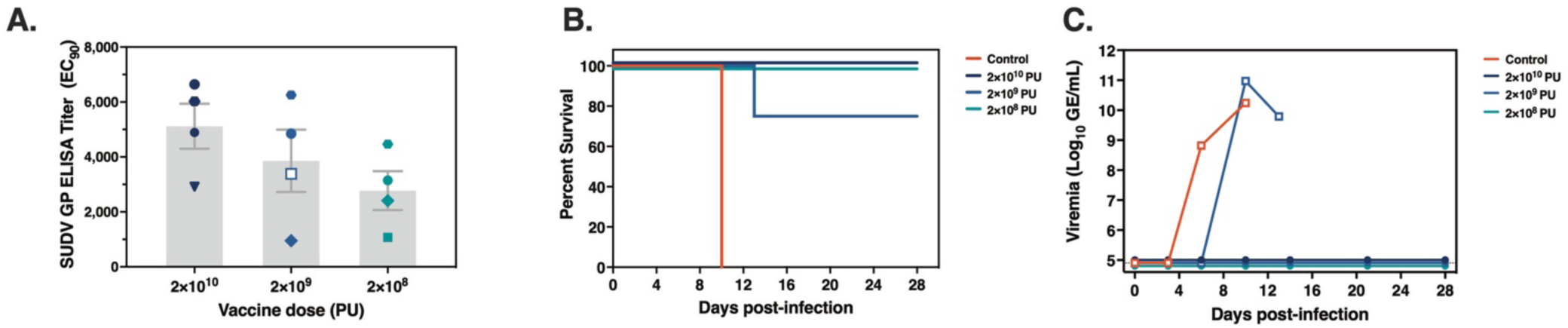
A single-shot bivalent ChAd3 vaccine protects from lethal SUDV challenge. Groups of macaques were immunized with a combination of ChAd3-EBOV and ChAd3-SUDV vaccine products at a 1:1 ratio (*e.g*., 1×10^10^ PU + 1×10^10^ PU) and challenged with 1,000 PFU SUDV/Bon at five weeks post-vaccination. (A) Anti-SUDV GP-specific ELISA titers at four weeks post-immunization (EC_90_). Open/white symbols indicate non-survivors. (B) Survival following lethal i.m. challenge with SUDV/Bon at five weeks post-immunization. (C) Plasma viremia by quantitative RT-PCR expressed as genomic equivalents per milliliter (GE/mL).

### A single-shot bivalent ChAd3-SUDV/ChAd3-EBOV vaccine protects against an aerosol challenge with SUDV/Bon and EBOV/Kik

Of particular concern for biodefense applications is protection from aerosolized threats (*27*). This has been a significant hurdle for vaccine development, as many vaccines that are successful in parenteral models of infection subsequently fail to protect against aerosol challenge. There are few reports of protection of macaques from aerosol challenge (*28–30*), leaving opportunity for effective vaccine development. To evaluate the efficacy of the single-shot, bivalent ChAd3 vaccine against aerosol challenge, groups of macaques were vaccinated with 1×10^10^ PU of ChAd3-SUDV + 1×10^10^ PU of ChAd3-EBOV and challenged 5 weeks later with either small particle aerosolized SUDV/Bon (Fig 5 A-C) or EBOV/Kik (Fig 5 D-E). Compared to unvaccinated control animals, the single-shot bivalent ChAd3 vaccine provided protection from both SUDV and EBOV aerosol infection, with only a single animal succumbing from either group (Fig 5 B and C). One control animal survived the aerosol SUDV exposure, which along with the single vaccinated death resulted in a non-significant difference between survival curves (Log-rank test *p* = 0.1). However, that animal had the lowest pre-challenge IgG titer (EC_90_ of 2888 at week 4 post-immunization, below the uniformly protective titer). Overall, the survival of vaccinated animals was promising with the bivalent vaccination resulting in a significant improvement of survival following EBOV aerosol exposure (*p* < 0.001) with transient viremia observed in a single surviving subject.

**Figure 5.**
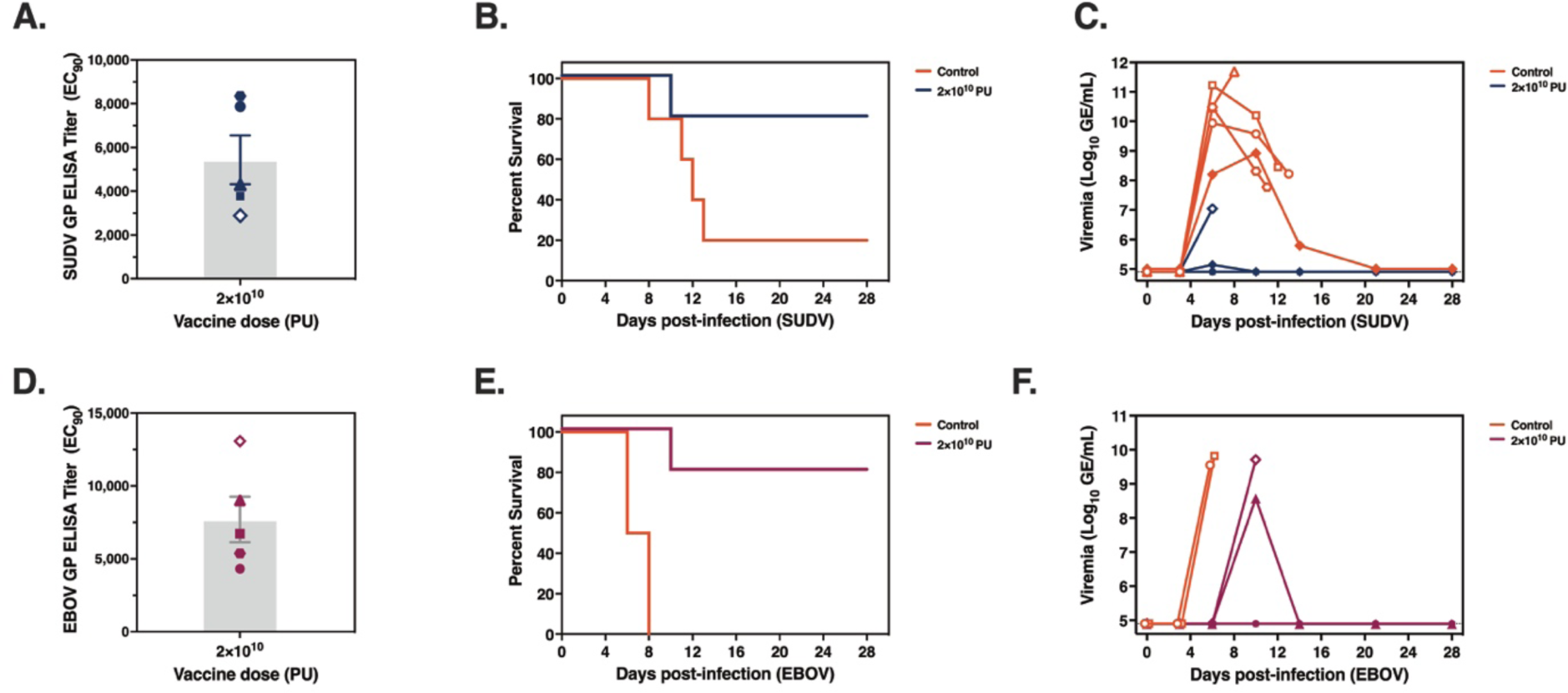
The bivalent ChAd3 vaccine protects animals from aerosol challenge. Groups of cynomolgus macaques (*n*=5) were either vaccinated with 1×10^10^ ChAd3-EBOV/ChAd3-SUDV or remained unvaccinated. (A) Anti-SUDV GP-specific ELISA titers at 4 weeks post-immunization. (B) Survival following aerosol challenge with SUDV/Bon at 5 weeks post-vaccination. (C) Plasma viremia by quantitative RT-PCR expressed as genomic equivalents per milliliter (GE/mL). (D) Anti-EBOV GP-specific ELISA titers at 4 weeks post-immunization. (E) Survival following aerosol challenge with EBOV/Kik at 5 weeks post-vaccination. (F) Plasma viremia by quantitative RT-PCR expressed as genomic equivalents per milliliter (GE/mL).

## DISCUSSION

Vaccines that are safe and effective with a simple regimen are essential for preparing for (re-)emerging viruses such as SUDV. EVD caused by SUDV is an epidemic disease with a high case fatality rate for which there is no licensed vaccine or therapeutics, points emphasized by the current outbreak in Uganda. Ideally, vaccines for epidemic viruses would induce both rapid and long-lasting protection, have a simple vaccine regimen for deployment in resource-poor environments and would utilize a vaccine platform with an established safety profile.

We have previously established that a single dose of a ChAd3-based vaccine targeting EBOV produced robust Ag-specific immune responses and was highly effective against lethal infection, protecting 100% of vaccinated macaques against acute challenge and providing one year of lasting immunity (*10*). Similarly, we have used this vector to demonstrate rapid and durable protection against MARV (*26*). For both vaccines, virus-specific IgG titers at acute timepoints were correlated with protection (*26*). The ChAd3 vaccine platform has been evaluated in multiple human clinical trials involving both adults and pediatrics, where it was found to be immunogenic and well tolerated (*11–14, 24*). Here, we demonstrate that a ChAd3-SUDV vaccine protected macaques from lethal challenge with SUDV by both i.m. and aerosol route of infection. We also defined anti-SUDV GP antibodies as the immunological correlate of protection, which will be essential for approval of a ChAd3-based SUDV vaccine by the FDA Animal Rule.

Protection from SUDV infection in the lethal NHP model by ChAd3-SUDV was rapid and only required a single shot. This is essential for vaccines used in areas with less developed public health infrastructure and immunization programs, since at-risk individuals may only have the opportunity for a single vaccination. The ideal SUDV vaccination regimen, in addition to protecting quickly following immunization, will generate a memory response sufficient for durable protection at extended timepoints. This will be critical for coverage of health care workers and community individuals throughout the duration of an outbreak. Here, a single dose of ChAd3-SUDV completely protected NHP challenged at 6 months and provided significant protection as late as 1 year with 3 of 4 NHP surviving. At six months, protective efficacy was 100% with a mean EC_90_ titer of 6390, a concentration which predicts 100% protection per the logistic regression shown in this study. At one-year, protective efficacy was 75% survival (3/4 NHP), where the animal that succumbed had an EC_90_ titer of 941 prior to challenge, a concentration where survival is not predicted. Interestingly, another animal with an EC_90_ titer of 328 survived SUDV challenge, indicating that the correlate of survival may not hold for the 12-month timepoint.

In the case where the immune correlate does not hold, it is intriguing to consider whether antibodies are a non-mechanistic immune correlate of protection, simply representing the overall vaccine-induced immune responses required for survival (*31*). In this case, protection observed may be explained by induction of an amnestic B-cell response upon challenge that leads to proliferation and differentiation of antigen-specific memory B cells resulting in protection at the memory time point or may also require T-cells. A mechanistic role for T lymphocytes in protection against EBOV infection has been demonstrated in animal models (*32–34*). However, previous studies by our group using ChAd3 for filovirus vaccines have not indicated a correlation between survival and T cell magnitude or quality, and incorporation of T cell responses did not strengthen the antibody correlate (*10*). Further, in our studies evaluating ChAd3-MARV, we showed uniform protection as soon as 1 week post-immunization, where there were no detectable antibodies, and did not find any correlation between neutralizing antibody titers and survival (*26*). These findings indicate that GP-specific binding titers are a robust, non-mechanistic correlate of protection that serve as a critical quantitative tool to bridge NHP efficacy to human immunogenicity data.

As outbreaks of filoviruses such as the current SUDV outbreak in Uganda are sporadic and unpredictable, often occurring in remote locations, vaccines for SUDV will likely require the FDA Animal Rule for licensure. In this alternative regulatory pathway, immunogenicity and safety data from human Phase 1/2 trials as well as efficacy and immunogenicity from NHP challenge studies will be required. Evaluation of the ChAd3-SUDV vaccine in a Phase I clinical trial (NCT04041570) assessing safety and immunogenicity in Uganda is complete and the vaccine has progressed toward advanced clinical development in Phase I/II clinical trials (*25*).

In conclusion, the data reported here demonstrate that ChAd3-SUDV vaccination elicits rapid and durable protection from lethal SUDV challenge. In conjunction with the positive safety profile reported in the Phase I clinical trial, makes it suitable for use as an outbreak-appropriate emergency vaccine to protect healthcare workers and affected communities.

## MATERIALS AND METHODS

### Animal vaccination and safety

Groups of 2–5 year old cynomolgus macaques (*Macaca fascicularis*) weighing between 2–3 kg were obtained from Covance for these immunization and challenge studies. All animal experiments were conducted under protocols approved by NIH and USAMRIID or UTMB Animal Care and Use committees. All experiments involving the use of SUDV or EBOV in animals were performed in a BSL-4 laboratory. The facilities used are fully accredited by the Association for Assessment and Accreditation of Laboratory Animal Care International. Cynomolgus macaque research was conducted in compliance with the Animal Welfare Act and other federal statutes and regulations relating to animals and experiments involving animals and adheres to the principles stated in the Guide for the Care and Use of Laboratory Animals, National Research Council.

The monkeys were singly housed singly and provided regular enrichment. Prior to blood sampling or vaccination, animals were anesthetized with ketamine or Telazol. Subjects received intramuscular vaccinations in the bilateral deltoids by needle and syringe with the doses and vectors indicated in figure legends and the text. Construction of the ChAd3 vaccines expressing SUDV or EBOV GP is detailed previously (*10*).

Following immunization, animals were transferred to the Maximum Containment Laboratory (BSL-4) for infection and remained there through study completion. The monkeys were fed and checked at least daily according to the protocol.

### Infectious challenge studies

For infectious virus challenges, either Sudan virus Boneface (SUDV/Bon) (*35, 36*), Sudan virus Gulu (SUDV/Gulu) (*37*) or Ebola virus Kikwit (EBOV/Kik) was used (*32, 38*). Macaques were challenged intramuscularly (I.M.) with a target dose of 1,000 PFU (plaque-forming units) of SUDV/Bon, SUDV/Gulu or EBOV/Kik in a volume of 1mL. For aerosol challenge, NHPs were exposed to a target dose of 100 PFU in a head-only chamber in a class III biological safety cabinet connected to the BSL-4 as previously described (*38, 39*). During the challenge studies, blood was collected from the NHPs for hematological, biochemical and virological analyses as previously described (*10, 32*). Following the development of clinical signs, animals were checked multiple times daily using institute-approved scoring criteria to determine timing of humane euthanasia under anesthesia to minimize pain and distress. Investigators were blinded to allocation during experiments and outcome assessment with un-blinding performed at the end of the challenge study.

### IgG ELISA

Methods for the GP IgG ELISA have been described previously (*10, 26, 40*). ELISA titers are expressed as EC_90_, reciprocal serum dilution values, which represent the dilution at which there is a 90% decrease in antigen binding.

### Statistical analyses

Statistical analyses were performed using GraphPad Prism version 9.4.1.

## Acknowledgments

This work was supported by the Intramural Research Program of the Vaccine Research Center, the National Institute of Allergy and Infectious Diseases, and the National Institutes of Health. We thank the staff of USAMRIID’s Viral Therapeutics Branch and Diagnostics Systems Division for technical contributions to the conduct of the EBOV/Kik i.m. study and the aerosol studies, as well as to the Aerobiology Division who provided support for the aerosol challenge. The authors also thank the UTMB Animal Resource Center for husbandry support of laboratory animals, Daniel Deer and Chad Mire for assistance with the animal studies, and Krystle Agans for performing PCR assays. Dr. John Dye provided the SUDV/Bon used for the aerosol challenge.

## Author conthributions

Conceptualization: NJS

Methodology: NJS

Formal analysis: ANH, RH, JIM, AP, DAS

Investigation: ANH, AP, CNM, LW, CC, WS, TM, DC, KF, JBG, TWG, JCT, JCJ, SEWR

Visualization: ANH

Project administration: ANH, TWG, NJS

Supervision: NJS

Writing – original draft: ANH, RH, JIM

Writing – review & editing: ANH, RH, JIM, NJS

## Data and materials availability

All data are available in the main text or the supplementary materials.

