## Supplementary files for "A Single-shot ChAd3 Vaccine Provides Protection from Intramuscular and Aerosol Sudan Virus Exposure"

### Supplementary Figures

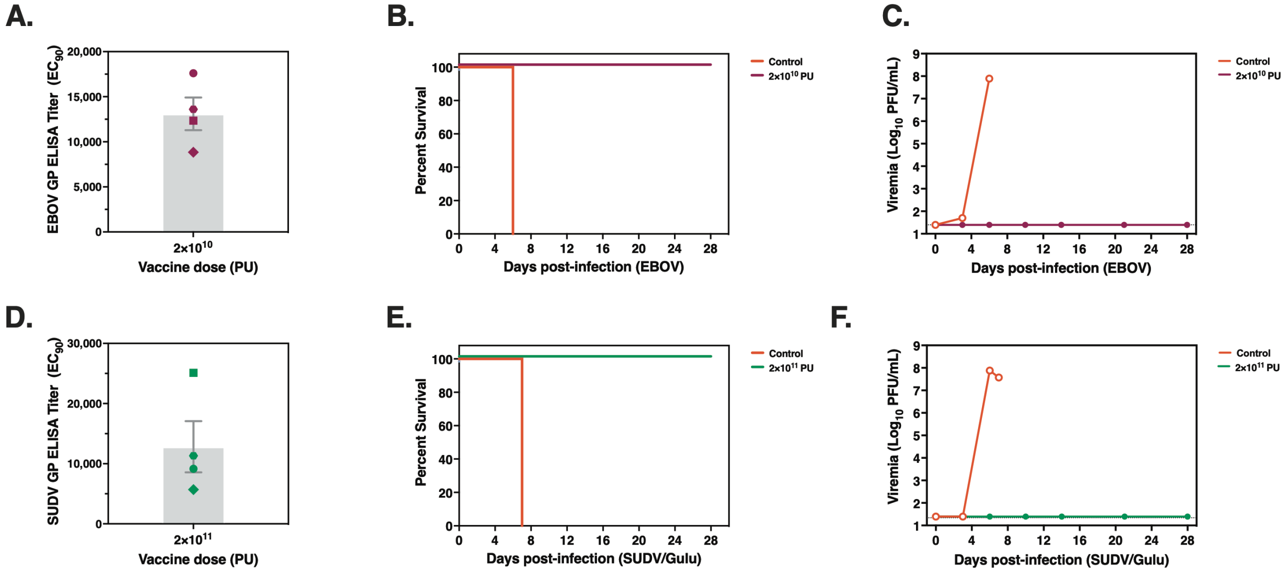


**Figure S1. A single-shot bivalent ChAd3 vaccine protects from lethal challenge with EBOV or SUDV/Gulu.** Groups of macaques were immunized with a combination of ChAd3-EBOV and ChAd3-SUDV vaccine products at a 1:1 ratio (e.g., $1\times10^{10}$ PU + $1\times10^{10}$ PU) and challenged with 1,000 PFU EBOV/Kik (A-C) or SUDV/Gulu (D-F) i.m. at five weeks post-vaccination. (A,D) GP-specific ELISA titers at four weeks post-immunization ($EC_{90}$). (B,E) Survival following lethal i.m. challenge at five weeks post-immunization. (C,F) Plasma viremia by plaque assay.

| | | | Immunization Phase | | | | | | | | | | | | |
|---|---|---|---|---|---|---|---|---|---|---|---|---|---|---|---|
| Figure | NHP ID | Vaccine | Dose | EC$_{90}$ titer (4 wk post-vaccine) | Virus | Route | Dose (PFU) | Interval from vaccine | Survived? | Day of death | Peak viremia (Log$_{10}$ PFU/mL or GE/mL) | BUN | ALT | AST | GGT |
| 1 & 2 | Ctl 1 | n/a | n/a | n/a | SUDV/Bon | IM | 650 | n/a | No | 10 | 6.54 | + | + | ++ | ++ |
| 1 & 2 | Ctl 2 | n/a | n/a | n/a | SUDV/Bon | IM | 1263 | n/a | No | 12 | 7.53 | +++ | +++ | +++ | + |
| 1 & 2 | NHP 1 | ChAd3-SUDV | 1.00E+10 | 3038 | SUDV/Bon | IM | 650 | 5 wk | Yes | 28 | < LOD | - | - | - | - |
| 1 & 2 | NHP 2 | ChAd3-SUDV | 1.00E+10 | 5897 | SUDV/Bon | IM | 1263 | 5 wk | Yes | 28 | < LOD | - | - | - | - |
| 1 & 2 | NHP 3 | ChAd3-SUDV | 1.00E+10 | 9295 | SUDV/Bon | IM | 1263 | 5 wk | Yes | 28 | < LOD | - | - | - | - |
| 1 & 2 | NHP 4 | ChAd3-SUDV | 1.00E+10 | 7659 | SUDV/Bon | IM | 1263 | 5 wk | Yes | 28 | < LOD | - | - | - | - |
| 1 & 2 | NHP 5 | ChAd3-SUDV | 1.00E+10 | 8661 | SUDV/Bon | IM | 1263 | 5 wk | Yes | 28 | < LOD | - | - | - | - |
| 2 | NHP 1 | ChAd3-SUDV | 1.00E+09 | 1712 | SUDV/Bon | IM | 650 | 5 wk | No | 8 | 8.19 | +++ | +++ | +++ | +++ |
| 2 | NHP 2 | ChAd3-SUDV | 1.00E+09 | 1608 | SUDV/Bon | IM | 1263 | 5 wk | No | 9 | 7.04 | - | - | - | - |
| 2 | NHP 3 | ChAd3-SUDV | 1.00E+09 | 1562 | SUDV/Bon | IM | 1263 | 5 wk | No | 10 | 7.53 | ++ | - | ++ | ++ |
| 2 | NHP 4 | ChAd3-SUDV | 1.00E+09 | 17303 | SUDV/Bon | IM | 1263 | 5 wk | Yes | 28 | < LOD | - | - | - | - |
| 2 | NHP 5 | ChAd3-SUDV | 1.00E+09 | 3648 | SUDV/Bon | IM | 1263 | 5 wk | Yes | 28 | < LOD | - | - | - | - |
| 2 | NHP 6 | ChAd3-SUDV | 1.00E+08 | 3278 | SUDV/Bon | IM | 650 | 5 wk | No | 9 | 8.09 | +++ | + | ++ | - |
| 2 | NHP 7 | ChAd3-SUDV | 1.00E+08 | 2371 | SUDV/Bon | IM | 1263 | 5 wk | No | 10 | 3.98 | ++ | ++ | +++ | ++ |
| 2 | NHP 8 | ChAd3-SUDV | 1.00E+08 | 1865 | SUDV/Bon | IM | 1263 | 5 wk | No | 9 | 5.70 | ++ | - | +++ | - |
| 2 | NHP 9 | ChAd3-SUDV | 1.00E+08 | 9460 | SUDV/Bon | IM | 1263 | 5 wk | Yes | 28 | < LOD | - | - | - | - |
| 2 | NHP 10 | ChAd3-SUDV | 1.00E+08 | 4120 | SUDV/Bon | IM | 1263 | 5 wk | Yes | 28 | < LOD | - | - | - | - |
| 2 | NHP 11 | ChAd3-SUDV | 1.00E+08 | 4169 | SUDV/Bon | IM | 1038 | 5 wk | Yes | 28 | < LOD | - | - | - | - |
| 2 | NHP 12 | ChAd3-SUDV | 1.00E+08 | 1219 | SUDV/Bon | IM | 1038 | 5 wk | No | 8 | 7.93 | ++ | +++ | +++ | ++ |
| 2 | NHP 13 | ChAd3-SUDV | 1.00E+08 | 13148 | SUDV/Bon | IM | 1038 | 5 wk | Yes | 28 | < LOD | - | - | - | - |
| 2 | NHP 14 | ChAd3-SUDV | 5.00E+07 | 387 | SUDV/Bon | IM | 1038 | 5 wk | No | 10 | 5.89 | +++ | +++ | +++ | + |
| 2 | NHP 15 | ChAd3-SUDV | 5.00E+07 | 1601 | SUDV/Bon | IM | 1038 | 5 wk | No | 11 | 5.8 | ++ | + | +++ | - |
| 2 | NHP 16 | ChAd3-SUDV | 5.00E+07 | 3621 | SUDV/Bon | IM | 1038 | 5 wk | Yes | 28 | < LOD | - | - | - | - |
| 2 | NHP 17 | ChAd3-SUDV | 1.00E+07 | 120 | SUDV/Bon | IM | 650 | 5 wk | No | 11 | 5.1 | +++ | + | +++ | ++ |
| 3 | Ctl 1 | n/a | n/a | n/a | SUDV/Bon | IM | 1350 | n/a | No | 9 | 7.94 | - | - | + | - |
| 3 | Ctl 2 | n/a | n/a | n/a | SUDV/Bon | IM | 1238 | n/a | No | 18 | 6.18 | - | + | + | - |
| 3 | NHP 1 | ChAd3-SUDV | 1.00E+11 | 10363 | SUDV/Bon | IM | 1350 | 6 mo | Yes | 28 | < LOD | - | - | - | - |
| 3 | NHP 2 | ChAd3-SUDV | 1.00E+11 | 17511 | SUDV/Bon | IM | 1350 | 6 mo | Yes | 28 | < LOD | - | - | - | - |
| 3 | NHP 3 | ChAd3-SUDV | 1.00E+11 | 10021 | SUDV/Bon | IM | 1350 | 6 mo | Yes | 28 | < LOD | - | - | - | - |
| 3 | NHP 4 | ChAd3-SUDV | 1.00E+11 | 290 | SUDV/Bon | IM | 1350 | 6 mo | Yes | 28 | < LOD | - | - | - | - |
| 3 | NHP 5 | ChAd3-SUDV | 1.00E+11 | 2868 | SUDV/Bon | IM | 1238 | 12 mo | Yes | 28 | 5.69 | - | - | - | - |
| 3 | NHP 6 | ChAd3-SUDV | 1.00E+11 | 13494 | SUDV/Bon | IM | 1238 | 12 mo | Yes | 28 | 5.58 | - | - | - | - |
| 3 | NHP 7 | ChAd3-SUDV | 1.00E+11 | 9197 | SUDV/Bon | IM | 1238 | 12 mo | Yes | 28 | < LOD | - | - | - | - |
| 3 | NHP 8 | ChAd3-SUDV | 1.00E+11 | 5505 | SUDV/Bon | IM | 1238 | 12 mo | No | 9 | 6.66 | ++ | + | +++ | ++ |

*Chemistry values elevation from baseline: BUN + 2-4 fold, ++ 4-7 fold, +++ >7 fold; ALT + 3-7 fold, ++ 7-15 fold, +++ >15 fold; AST + 4-10 fold, ++ 10-20 fold, +++ >20 fold; GGT + 2-3 fold, ++ 3-6 fold, +++ >6 fold*

| | | Immunization Phase | | | Challenge Phase | | | | | | | | | | |
|---|---|---|---|---|---|---|---|---|---|---|---|---|---|---|---|
| Figure | NHP ID | Vaccine | Dose | $EC_{90}$ titer (4 wk post-vaccine) | Virus | Route | Dose (PFU) | Interval from vaccine | Survived? | Day of death | Peak viremia ($Log_{10}$ PFU/mL or GE/mL) | BUN | ALT | AST | GGT |
| 4 | Ctl 1 | n/a | n/a | n/a | SUDV/Bon | IM | 1000 | n/a | No | 10 | 10.24 | +++ | ++ | +++ | ++ |
| 4 | NHP 1 | ChAd3-SUDV + ChAd3-EBOV | 2.00E+10 | 6642 | SUDV/Bon | IM | 1000 | 5 wk | Yes | 28 | < LOD | - | - | - | - |
| 4 | NHP 2 | ChAd3-SUDV + ChAd3-EBOV | 2.00E+10 | 6024 | SUDV/Bon | IM | 1000 | 5 wk | Yes | 28 | < LOD | - | - | - | - |
| 4 | NHP 3 | ChAd3-SUDV + ChAd3-EBOV | 2.00E+10 | 2913 | SUDV/Bon | IM | 1000 | 5 wk | Yes | 28 | < LOD | - | - | - | - |
| 4 | NHP 4 | ChAd3-SUDV + ChAd3-EBOV | 2.00E+10 | 4891 | SUDV/Bon | IM | 1000 | 5 wk | Yes | 28 | < LOD | - | - | - | - |
| 4 | NHP 5 | ChAd3-SUDV + ChAd3-EBOV | 2.00E+09 | 950 | SUDV/Bon | IM | 1000 | 5 wk | Yes | 28 | < LOD | - | - | - | - |
| 4 | NHP 6 | ChAd3-SUDV + ChAd3-EBOV | 2.00E+09 | 3383 | SUDV/Bon | IM | 1000 | 5 wk | No | 13 | 10.97 | +++ | ++ | +++ | +++ |
| 4 | NHP 7 | ChAd3-SUDV + ChAd3-EBOV | 2.00E+09 | 4848 | SUDV/Bon | IM | 1000 | 5 wk | Yes | 28 | < LOD | - | - | - | - |
| 4 | NHP 8 | ChAd3-SUDV + ChAd3-EBOV | 2.00E+09 | 6255 | SUDV/Bon | IM | 1000 | 5 wk | Yes | 28 | < LOD | - | - | - | - |
| 4 | NHP 9 | ChAd3-SUDV + ChAd3-EBOV | 2.00E+08 | 1073 | SUDV/Bon | IM | 1000 | 5 wk | Yes | 28 | < LOD | + | - | - | - |
| 4 | NHP 10 | ChAd3-SUDV + ChAd3-EBOV | 2.00E+08 | 3148 | SUDV/Bon | IM | 1000 | 5 wk | Yes | 28 | < LOD | - | - | - | - |
| 4 | NHP 11 | ChAd3-SUDV + ChAd3-EBOV | 2.00E+08 | 4465 | SUDV/Bon | IM | 1000 | 5 wk | Yes | 28 | < LOD | - | - | - | - |
| 4 | NHP 12 | ChAd3-SUDV + ChAd3-EBOV | 2.00E+08 | 2410 | SUDV/Bon | IM | 1000 | 5 wk | Yes | 28 | < LOD | - | - | - | - |
| 5 | Ctl 1S | n/a | n/a | n/a | SUDV/Bon | Aero | 100 * | n/a | No | 13 | 9.94 | +++ | + | +++ | + |
| 5 | Ctl 2S | n/a | n/a | n/a | SUDV/Bon | Aero | 100 * | n/a | No | 12 | 11.22 | +++ | ++ | ++ | ++ |
| 5 | Clt 3S | n/a | n/a | n/a | SUDV/Bon | Aero | 100 * | n/a | Yes | 28 | 8.82 | - | - | - | - |
| 5 | Ctl 4S | n/a | n/a | n/a | SUDV/Bon | Aero | 100 * | n/a | No | 11 | < LOD | +++ | - | - | ++ |
| 5 | Ctl 5S | n/a | n/a | n/a | SUDV/Bon | Aero | 100 * | n/a | No | 8 | 11.68 | ++ | + | ++ | ++ |
| 5 | NHP 1S | ChAd3-SUDV + ChAd3-EBOV | 2.00E+10 | 7872 | SUDV/Bon | Aero | 100 * | 5 wk | Yes | 28 | < LOD | - | - | - | - |
| 5 | NHP 2S | ChAd3-SUDV + ChAd3-EBOV | 2.00E+10 | 3777 | SUDV/Bon | Aero | 100 * | 5 wk | Yes | 28 | 10.48 | - | - | - | - |
| 5 | NHP 3S | ChAd3-SUDV + ChAd3-EBOV | 2.00E+10 | 2888 | SUDV/Bon | Aero | 100 * | 5 wk | No | 10 | 7.04 | - | - | - | - |
| 5 | NHP 4S | ChAd3-SUDV + ChAd3-EBOV | 2.00E+10 | 8349 | SUDV/Bon | Aero | 100 * | 5 wk | Yes | 28 | < LOD | - | - | - | - |
| 5 | NHP 5S | ChAd3-SUDV + ChAd3-EBOV | 2.00E+10 | 4307 | SUDV/Bon | Aero | 100 * | 5 wk | Yes | 28 | < LOD | - | - | - | - |
| 5 | Ctl 1E | n/a | n/a | n/a | EBOV/Kik | Aero | 100 * | n/a | No | 8 | 9.82 | - | - | + | - |
| 5 | Ctl 2E | n/a | n/a | n/a | EBOV/Kik | Aero | 100 * | n/a | No | 8 | 9.55 | - | - | - | + |
| 5 | NHP 1E | ChAd3-SUDV + ChAd3-EBOV | 2.00E+10 | 4318 | EBOV/Kik | Aero | 100 * | 5 wk | Yes | 28 | < LOD | - | - | - | - |
| 5 | NHP 2E | ChAd3-SUDV + ChAd3-EBOV | 2.00E+10 | 13076 | EBOV/Kik | Aero | 100 * | 5 wk | No | 10 | 9.71 | ++ | +++ | +++ | +++ |
| 5 | NHP 3E | ChAd3-SUDV + ChAd3-EBOV | 2.00E+10 | 9025 | EBOV/Kik | Aero | 100 * | 5 wk | Yes | 28 | < LOD | - | - | - | - |
| 5 | NHP 4E | ChAd3-SUDV + ChAd3-EBOV | 2.00E+10 | 5376 | EBOV/Kik | Aero | 100 * | 5 wk | Yes | 28 | 8.58 | - | - | - | - |
| 5 | NHP 5E | ChAd3-SUDV + ChAd3-EBOV | 2.00E+10 | 6725 | EBOV/Kik | Aero | 100 * | 5 wk | Yes | 28 | < LOD | - | - | - | - |
| S1 | Ctl 1E | n/a | n/a | n/a | EBOV/Kik | IM | 1275 | n/a | No | 6 | 7.89 | ++ | ++ | +++ | ++ |
| S1 | NHP 1E | ChAd3-SUDV + ChAd3-EBOV | 2.00E+10 | 4971 | EBOV/Kik | IM | 1275 | 5 wk | Yes | 28 | < LOD | - | - | - | - |
| S1 | NHP 2E | ChAd3-SUDV + ChAd3-EBOV | 2.00E+10 | 8416 | EBOV/Kik | IM | 1275 | 5 wk | Yes | 28 | < LOD | - | - | - | - |
| S1 | NHP 3E | ChAd3-SUDV + ChAd3-EBOV | 2.00E+10 | 9075 | EBOV/Kik | IM | 1275 | 5 wk | Yes | 28 | < LOD | - | - | - | - |
| S1 | NHP 4E | ChAd3-SUDV + ChAd3-EBOV | 2.00E+10 | 10328 | EBOV/Kik | IM | 1275 | 5 wk | Yes | 28 | < LOD | - | - | - | - |
| S1 | Ctl 1S | n/a | n/a | n/a | SUDV/Gulu | IM | 875 | 5 wk | No | 7 | 7.88 | - | ++ | +++ | ++ |

| | | | | | | | | | | | | | | |
|---|---|---|---|---|---|---|---|---|---|---|---|---|---|---|
| S1 | NHP 5S | ChAd3-SUDV + ChAd3-EBOV | 2.00E+11 | SUDV/Gulu | IM | 875 | 5 wk | Yes | 28 | < LOD | - | - | - | - |
| S1 | NHP 6S | ChAd3-SUDV + ChAd3-EBOV | 2.00E+11 | SUDV/Gulu | IM | 875 | 5 wk | Yes | 28 | < LOD | - | - | - | - |
| S1 | NHP 7S | ChAd3-SUDV + ChAd3-EBOV | 2.00E+11 | SUDV/Gulu | IM | 875 | 5 wk | Yes | 28 | < LOD | - | - | - | - |
| S1 | NHP 8S | ChAd3-SUDV + ChAd3-EBOV | 2.00E+11 | SUDV/Gulu | IM | 875 | 5 wk | Yes | 28 | < LOD | - | - | - | - |

*Chemistry values elevation from baseline: BUN + 2-4 fold, ++ 4-7 fold, +++ >7 fold; ALT + 3-7 fold, ++ 7-15 fold, +++ >15 fold; AST + 4-10 fold, ++ 10-20 fold, +++ >20 fold; GGT + 2-3 fold, ++ 3-6 fold, +++ >6 fold* ** Aerosol target dose was 100 PFU/NHP*
